# Fungal auxin is a quorum-based modulator of blast disease severity

**DOI:** 10.1101/2021.10.26.465851

**Authors:** Lihong Dong, Qing Shen, Cheng-yen Chen, Lizheng Shen, Fan Yang, Naweed I. Naqvi, Yi Zhen Deng

## Abstract

Auxin is an important phytohormone regulating plant growth and development, and can also be produced by microbial pathogens including the rice-blast fungus *Magnaporthe oryzae*. However, the detailed biosynthesis pathway, biological function(s), and cellular distribution of such fungal auxin in *M. oryzae* remain largely unknown. Here, we report a sequential accumulation of intrinsic auxin in the three conidial cells, the infection structure (appressorium), and the invasive hyphae in *M. oryzae*. Such fungus-derived auxin was also secreted out and perceived by the host plants. A mitochondria-associated Indole-3-pyruvate decarboxylase, Ipd1, is essential for auxin/Indole-3-acetic acid biosynthesis in *M. oryzae*. The *ipd1*Δ was defective in pathogenicity whereas overexpression of *IPD1* led to enhanced virulence in rice. Chemical inhibition of fungal IAA biosynthesis, or its increase via external supplementation decreased or increased the severity of blast disease, respectively, in a dose-dependent manner. Furthermore, the IAA produced and secreted by *M. oryzae* governed the incidence and severity of blast disease in a quorum-dependent manner. Appressorium formation, conidial cell death critical for appressorium function, and the transcription of infection-related genes, *MPG1* and *INV1*, directly correlated with cell density and/or IAA levels within the conidial population at the early stages of pathogenic development. Overall, our study revealed that the severity of blast disease is regulated via quorum sensing with intrinsic IAA serving as an associated signal transducer in rice blast.

## Introduction

Quorum sensing (QS), initially identified in bacteria, has now been established as an important regulatory mechanism of gene expression on a cell-density basis [1]. In a given group, an individual bacterium can secrete and perceive multiple chemical moieties called quorum-sensing molecules (QSMs), which increase in concentration as a function of cell density [2–3]. When the concentration of QSMs reaches a threshold, indicating that the bacterial population is sufficiently large, the individual cells in the group begin to simultaneously express genes governing virulence, antibiotic resistance, biofilm formation, and host immunity suppression or evasion, and therefore coordinate bacterial pathogenicity as a group [2–3]. Recent studies identified several QSMs in fungi, including *Candida albicans, Aspergillus flavus, Neurospora crassa, Penicillium sclerotiorum*, and *Alternaria crassa* [4–8]. In *Magnaporthe oryzae*, a secreted invertase (Inv1) is suggested as a potential QSM catalyzing the hydrolysis of extracellular sucrose to glucose and fructose, thus facilitating better nutrient acquisition during pathogenesis [9]. However, whether quorum-sensing mechanism indeed contributes to plant infection by the rice blast fungus is unclear, and whether phytohormones can also mediate quorum-sensing behavior is unknown.

Leaf and panicle blast is a serious disease destroying rice, wheat and other cereals each year that are more than enough to feed 60 million people [10–12]. It is caused by the filamentous ascomycete *M. oryzae*, which produces asexual spores known as conidia that can germinate and form a special cell named an appressorium, at the tip of the germ tube, for host infection [10, 13-14]. Phytohormones are known to mediate chemical communication between *M. oryzae* and host rice [15]. It has been reported that the accumulation of auxin/Indole-3-acetic acid (IAA) in rice causes enhanced susceptibility to *M. oryzae*, while blocking rice IAA synthesis or signaling in rice helps gain resistance to blast disease [16]. *M. oryzae*, has also been reported to produce IAA in its vegetative hyphae and conidia [17]. However, it remains unclear about the physiological function of such fungus-derived auxin/IAA and its contribution to the establishment of the blast disease.

In this study, we found intracellular auxin/IAA, synthesized by a mitochondria-associated pyruvate decarboxylase named as Ipd1, accumulated in a sequential manner starting from terminal conidial cell that will undergo cell death subsequently. More importantly, we found that *M. oryzae* infects the host in an auxin-level and/or cell-density dependent manner. The fungus-derived auxin/IAA was secreted out and determined conidial cell death, apressorium function, and transcription of the infection-related genes through quorum sensing. Our data provide insight into the hitherto unknown role of such fungus-derived phytohormone auxin/IAA as a novel QSM that regulates pathogenicity in *M. oryzae*.

## Results

### Intrinsic auxin accumulation in *M. oryzae* during pathogenic development and host infection

To investigate the homeostasis of endogenous auxin in *M. oryzae*, we constructed a fluorescent reporter strain by expressing the Domain II (DII) of plant auxin repressor protein [18] fused to Venus and a nuclear localization signal (NLS) [19] under the control of *RP27* promoter (Figure S1A-C). A histone H1-mCherry fusion protein was simultaneously co-expressed to mark the nuclei. Two independent strains co-expressing DII-Venus-NLS and H1-mCherry (DII-Venus for short) (#16 and #17) were analyzed and found to be indistinguishable from the wild type in terms of vegetative growth (Figure S1D and Table S1), conidiation (Table S1; p>0.05 vs. wild type), or pathogenesis (Figure S1E). The DII-Venus epifluorescence is expected to decrease/disappear upon auxin accumulation [19]. To verify whether DII-Venus indeed functions as an auxin reporter in *M. oryzae*, we treated the mycelia (strain #16 as a representative) with IAA, and found that the nuclear accumulation of the DII-Venus signal was abolished, whereas the H1-mCherry signal remained intact (Figure S2), thus proving that the DII-Venus was responsive to elevated level of auxin/IAA, and could reflect the endogenous auxin/IAA fluctuations in *M. oryzae*.

After validating the engineered DII-Venus as an auxin biosensor in *M. oryzae*, we then proceeded to monitor the endogenous auxin homeostasis during pathogenic development and invasive growth. We were interested to first address whether auxin synthesis and/or distribution is uniform in the three conidial cells, which undergo regulated cell death one by one sequentially [20]. Interestingly, such intrinsic (fungal) auxin first accumulated in the terminal cell (the conidial cell distal to appressorium) at 8 hours post inoculation (hpi), and such auxin accumulation could be suppressed by the specific inhibitor amino-oxyacetic acid (AOA) (Figure 1A). At the late stage (24 hpi) when *M. oryzae* is ready to initiate host penetration, the auxin accumulation was found to be significantly higher in the appressorium (Figure 1A), thus implying that endogenous auxin plays a role in appressorium-mediated host penetration and/or invasive growth. Likewise, the fungal auxin showed enhanced accumulation in the invasive hyphae in rice sheath at the later stages of infection (48 hpi) as compared to the early host penetration phase (24 hpi), based on the changes of DII-Venus signal intensity during such invasive growth (Figure 1B). Our results showed that endogenous auxin sequentially accumulates in the developing conidia and appressoria, as well as in the invasive hyphae in *M. oryzae* during establishment of the blast disease in rice.

**Figure 1.**
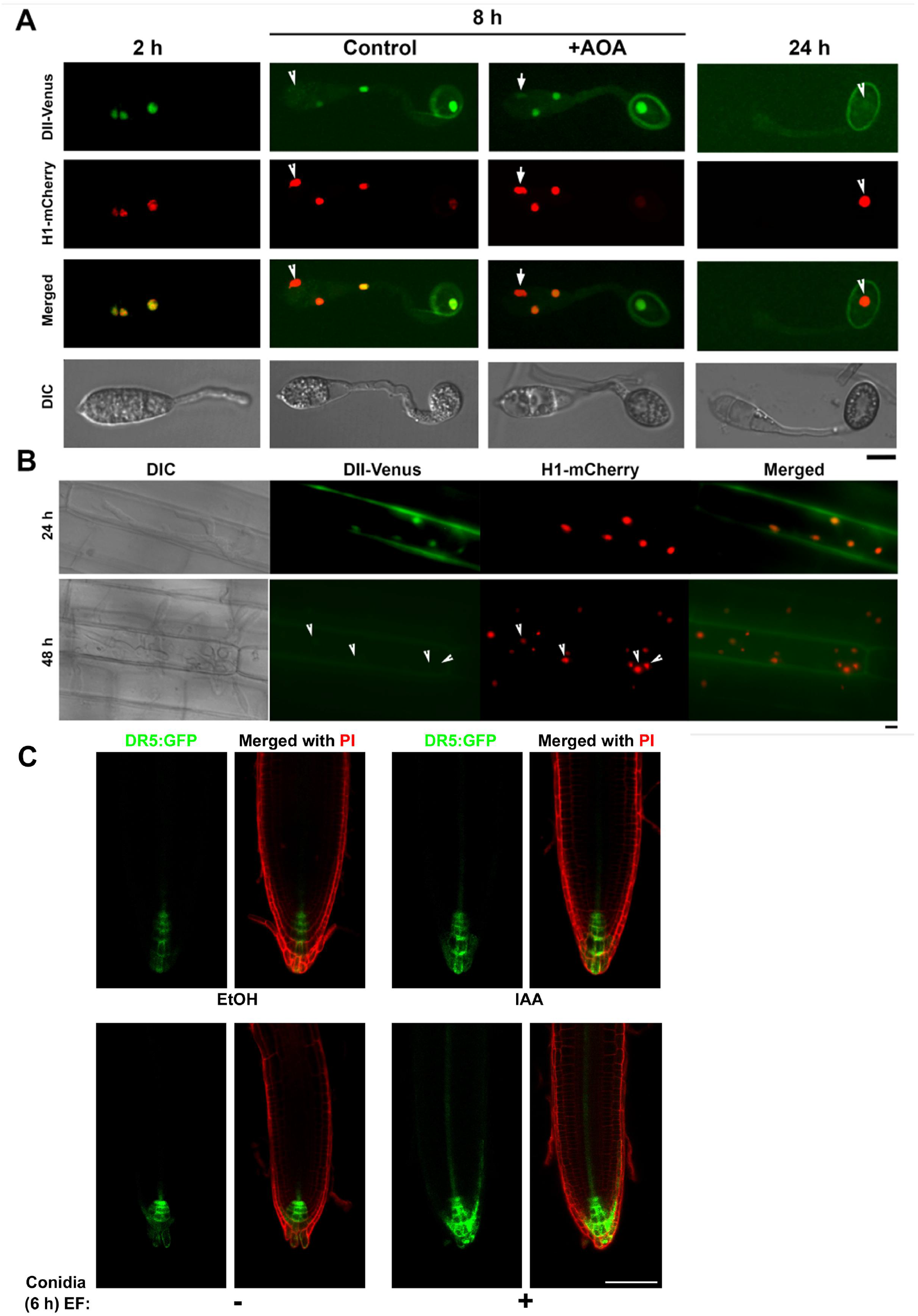
Intrinsic auxin accumulates in *M.oryzae*, and is secreted out and sensed by plant. (A) Auxin accumulation, reversely shown by the disappearance of DII-Venus signal, in the conidium and appressorium. Conidia of DII-Venus: H1-mCherry co-expressing strain (10^5^ /mL) were inoculated on inductive surface (cover slips) for appressorium formation and imaged at indicated time points. AOA (1 mM) was added at 4 hours post inoculation (hpi) to conidia that already formed appressoria. Arrowheads denote the nuclei (visualized by H1-mCherry) that have no DII-Venus signal, and arrows mark the nucleus with weak DII-Venus signal under AOA inhibition. Scale bar = 5 μm. (B) Auxin in the penetration and invasive hyphae, at 24 and 48 hpi, respectively. Conidia (10^5^ /mL) were inoculated on rice leaf sheath for *in planta* observation. Arrowheads mark the disappearance of DII-Venus signal. Bar =10 μm. (C) Secreted fungal auxin perceived by plant. The extracellular fluid (EF) with or without *M. oryzae* conidia (10^6^ /mL) was collected at 6 hpi and used to incubate with the seven-day-old DR5:GFP seedlings (Arabidopsis) for 30 min, before propidium iodide (PI) staining and epifluorescent microscopy. IAA (50 μM) served as the positive control, and ethanol (EtOH, 0.1%) as the solvent control of IAA. Scale bar = 100 μm. The DII-Venus and H1-mCherry signals were observed using a spinning disk confocal microscope, while DR5 and PI were observed using a Leica TCS SP8 X inverted microscope system. Images shown in this figure are all single plane images.

Next, we collected the extracellular fluid from *M. oryzae* conidia inoculated on the inductive surface for 6 hour (h), and incubated it with the *Arabidopsis* root expressing the DR5:GFP as an auxin biosensor (Figure 1C) [21]. The DR5:GFP signal was specifically induced by IAA or the extracellular fluid from the developing conidia, but not by the blank or solvent control (Figure 1C), demonstrating that such fungus-derived auxin could be secreted out and sensed by plant.

Overall, we conclude that *M. oryzae* produces and secretes auxin/IAA during pathogenic development. We infer that such fungal auxin/IAA is important for fungal pathogenesis as well as biotic interactions in the rice blast pathosystem.

### Auxin biosynthesis is catalyzed by the pyruvate decarboxylase Ipd1 in *M. oryzae*

We next sought to uncover the biosynthesis pathway of auxin in *M. oryzae*. Microbes (including fungi) could produce IAA through tryptophan-dependent pathway by deaminating tryptophan to Indole-3-Pyruvic Acid (IPA), which is decarboxylated to produce Indole-3-acetaldehyde (IAAld), a direct precursor of IAA production [22–23]. An Indole-3 pyruvate decarboxylase (IPDC) enzyme, LmIPDC2, is involved in auxin biosynthesis in the phytopathogenic fungus *Leptosphaeria maculans*, likely via catalyzing the IPA decarboxylation [24]. By BLAST (Basic Local Alignment Search Tool, https://blast.ncbi.nlm.nih.gov/Blast.cgi) search with LmIPDC2 (XP_003844157.1) as the bait, we identified an *M. oryzae* ortholog (MGG_01892), hereafter named as Ipd1. Phylogenetic analysis among yeast and filamentous fungi showed that Ipd1 is closer to ascomycetous fungi rather than yeast or basidiomycetous fungi (Figure S3A). To investigate possible function of Ipd1 in auxin/IAA production and/or *M. oryzae* pathogenicity, we generated a gene deletion mutant (Figure S3B-C), a genetic complementation strain (Figure S3D), and an overexpression strain (Figure S3E-F) of *IPD1*.

Using comparative liquid chromatography mass spectrometry (LC-MS) with IAA as a standard, we detected both the intracellular and extracellular (secreted) auxin/IAA in the wild-type mycelia in *M. oryzae* (Figure S4), which is consistent with what we saw during its pathogenic development (Figure 1A and C). The peak with the m/z ratio of 129.8, and molecular weight of 175.18 corresponded to the IAA metabolite (Figure S4-5). Such IAA accumulation in the *ipd1*Δ mutant or extracellular fluid (secreted) was reduced to a level of around 7% or 14% of that of the wild-type mycelia, respectively (Figure S4). In contrast, the intracellular IAA level showed nearly two-fold increase in the *IPD1* over-expression (*IPD1*-OX) strain as compared to the wild type, while the extracellular IAA level of the *IPD1*-OX strain was comparable to the wild type (Figure S4). We further found that treatment with the IAA biosynthesis inhibitor AOA mimicked the *IPD1* deletion phenotype, and resulted in significant reduction of both intracellular (22% of untreated control) and extracellular (12% of untreated control) IAA in *M. oryzae* (Figure S5). Overall, our results demonstrate that the IPDC enzyme Ipd1 is involved in IAA production, and thus could be used for assessing fungal auxin/IAA function in *M. oryzae* pathogenicity.

### Fungus-derived auxin determines the successful establishment of blast disease

Next we investigated the contribution of Ipd1-mediated auxin synthesis to *M. oryzae* infection ability. We were interested to notice that, the *IPD1*-OX strain, which produces a higher level of IAA (Figure S4) as compared with wild type, was able to cause advanced onset and severity of blast lesion on rice plants (Figure 2A). On the other hand, decrease IAA production by treating the wild-type conidial suspension with the auxin inhibitor AOA (Figure 1A, S5) or through *IPD1* deletion (Figure S4), lead to substantial reduction or complete loss of blast lesion in a dose-dependent manner (Figure 2A, S6A). Furthermore, AOA was less effective to restrict blast disease when applied to the *IPD1*-OX strain since a higher level of AOA was required to block the lesion formation as compared to the wild type (Figure 2A), indicating that elevated level of fungal IAA promotes rice infection. To confirm this, we treated the wild-type conidia with AOA and IAA, individually or in combination. More importantly, both chemicals were removed before *M. oryzae* starts host penetration, that is immediately after the infection-related morphogenesis/development (0-22 hpi), to minimize the effect of these chemicals on the rice host. We found that IAA-treated *M. oryzae* was more virulent, whereas AOA inhibition on rice blast fungus resulted in almost loss of such virulence, which could not be reverted with exogenous IAA (Figure 2B, left). Furthermore, the AOA-based inhibition of *M. oryzae* pathogenicity remained unchanged even if the chemical was not removed at 22 hpi (Figure 2B, right), suggesting that the reduction of fungal IAA levels, caused by AOA inhibition, is more likely responsible for the suppression of rice blast. Interestingly, we noticed that when conidial density/load decreases from 2000 to 125 conidia/droplet, *M. oryzae* pathogenicity reduced concomitantly, as indicated by lesion development on the same rice leaf, and such reduction of pathogenicity was abolished when IAA was supplied to the low density inoculum (125 conidia/droplet, Figure 2C). In addition, keeping the conidial density constant, IAA was able to promote blast lesion development in a dose-dependent manner (Figure 2D). Together, these data demonstrate that fungus-derived auxin determines the successful establishment of blast disease in a dose-dependent manner.

**Figure 2.**
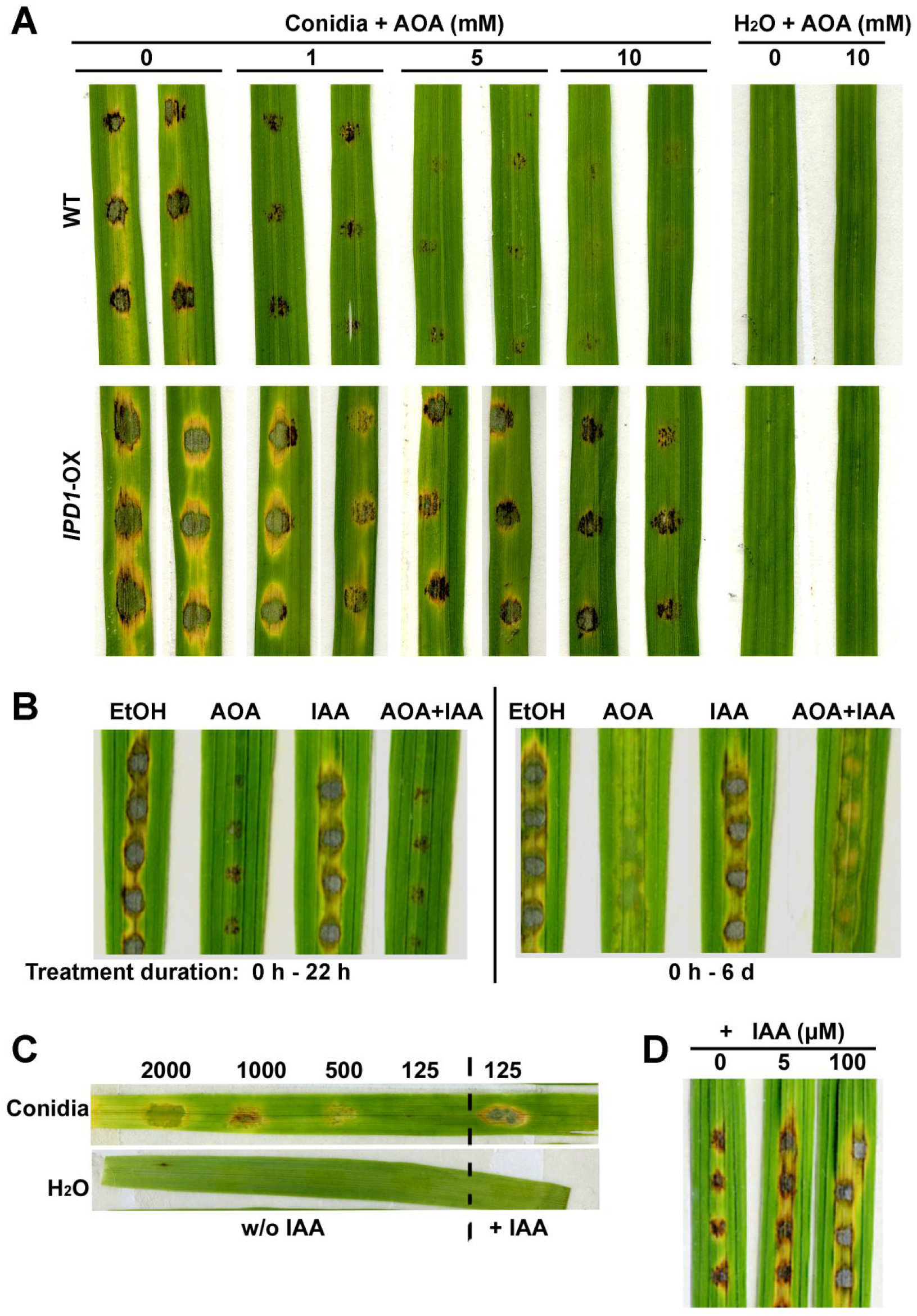
Ipd1-mediated auxin production determines successful establishment of blast disease in a dose-dependent manner. (A) Blast lesion caused by wild type or *IPD1* overexpression strain under different doses of AOA inhibition. Conidial suspension of wild-type (WT) or *IPD1*-OX strains, containing AOA of different concentrations, was inoculated on rice leaf explants. AOA was dissolved in sterile water (pH=7.0). Rice leaf explants inoculated with H2O, with or without AOA, served as blank controls. (B) Rice blast suppression caused by AOA inhibition of fungal IAA synthesis. Wild-type conidial suspension with AOA and/or IAA was inoculated on rice leaf explants. 0.02% ethanol (EtOH, solvent control of IAA), AOA (20 mM), and IAA (10 μM) were added into the conidial suspension at 0 hpi (h), and washed off at 22 h (left panel) or remained for 6 days (d) before photographing (right panel). (C) Blast lesion development on the basis of conidial cell density/IAA level. The conidial suspensions of serial dilution without (w/o) auxin supplementation were inoculated on the same rice leaf explant (left side of the dashed line). IAA (5 μM) was added to (+) the other repeat of inoculum with lowest cell density (125 conidia/mL, right side of the dashed line). Inoculation with water droplets, with or without IAA, served as negative controls. Photos were taken at 7 dpi. (D) The ability of IAA to promote *M.oryzae* pathogenicity dose-dependently, with constant conidial cell density. The wild-type conidial suspension supplemented with different concentrations of IAA was inoculated on the rice leaf explants, and photos were taken at 7 dpi.

### Fungal auxin acts as a quorum-sensing molecule regulating pathogenic development

Given the dosage-related behavior/phenotypes and the fact that fungal auxin is secreted out during pathogenic development, we hypothesized that such fungus-derived IAA could serve a quorum-sensing function that accumulates in a cell-density dependent manner while regulating pathogenic development in *M. oryzae*. To test this hypothesis, we first examined the effect of cell density on conidial cell death during appressorium development, since we saw a close correlation between fungal auxin accumulation and ferroptotic cell death in the conidium, which is critical for appressorium function [20, 25-26]. At 15 hpi, the condia from a low-density group (0.5×10^5^) displayed obviously longer germ tube, and delayed conidial cell death, as compared to the conidia from a high-density group (3×10^5^; Figure 3A). Adding IAA (100 μM) to the low-density group of conidia could effectively shorten the germ tube length and promote conidial cell death to a high-density level, while in contrast, addition of AOA (1 mM) to the high-density group of conidia led to long germ tube and delayed conidial cell death, mimicking those from low-density group (Figure 3A). We also noticed that the number of conidia able to form appressorium decreased as cell density declined and therefore systematically quantify the rate at 8 hpi. Indeed, percentage of conidia able to form appressorium decreased when the number of conidia in a given group reduced (Figure 3B). Again, such density dependent decrease of appressorium formation rate can be reversed by IAA supplementation (Figure 3B), whereas AOA completely blocked appressorium formation though the conidial density is high (Figure 3B). Interestingly, such AOA effect on appressorium formation turned out to be dose-dependent, for higher dosage blocked appressorium formation while lower dosage only delayed its formation (Figure S6B). Along with appressorium formation, we also quantified the cell death of the terminal conidial cell (Figure 3B) for intrinsic fungal IAA accumulate dominantly in this particular dying cell at 8 hpi (Figure 1A). Same experiment was perform to quantify conidial cell death except that AOA was added at 4 h to avoid its negative effect on appressorium formation. Similarly, number of conidia able to undergo cell death in the terminal cell declined along with cell density or IAA levels, whereas external IAA rescued the cell death defect of low-density conidia to a similar level of those from high-density group (Figure 3B). We further went on to test the effect of intrinsic IAA on conidial cell death at 24 hpi, when the entire conidium finished cell death, using different concentrations of auxin inhibitor and constant cell density. Similar to its effect on appressorium formation, AOA inhibited conidial cell death dose-dependently (Figure 3C). What’s more, we found that another well-established auxin inhibitor, L-kyrurenine (L-Kyn) [27], also effectively suppressed conidial death as AOA did (Figure 3C), confirming that it was the block of fungal auxin/IAA synthesis that led to suppression of conidial cell death. Based on these results, we infer that *M. oryzae* regulates pathogenic development (appressorium formation and conidial cell death) on a quorum-sensing basis using IAA as a quorum-sensing molecule (QSM), which is further supported by the close correlation between conidial density/IAA levels and length of germ tube when appressorium is formed (Figure 3D). Under low cell density, the conidia tended to delay appressorium formation and thus developed longer germ tubes (Figure 3A), to an average of approximately 4 μm at 8 hpi (Figure 3D), which may also contribute to the reduced/loss of pathogenicity in the low cell-density inoculum of conidia on the rice leaf (Figure 2C). However, addition of exogenous IAA reduced germ tube growth to a comparable level, average of nearly 2.5 μm at 8 hpi, as seen in (untreated) higher cell-density inoculum (Figure 3 A and D). Moreover, overexpression of *IPD1* resulted in reduced germ tube length (2-2.5 μm) even at a low cell-density inoculum, mimicking the effect of addition of IAA to the wild-type conidia at low cell-density (Figure 3D). All together, these results confirmed that pathogenic development of *M.oryzae* is regulated by the cell density-dependent Auxin/IAA accumulation through quorum sensing.

**Figure 3.**
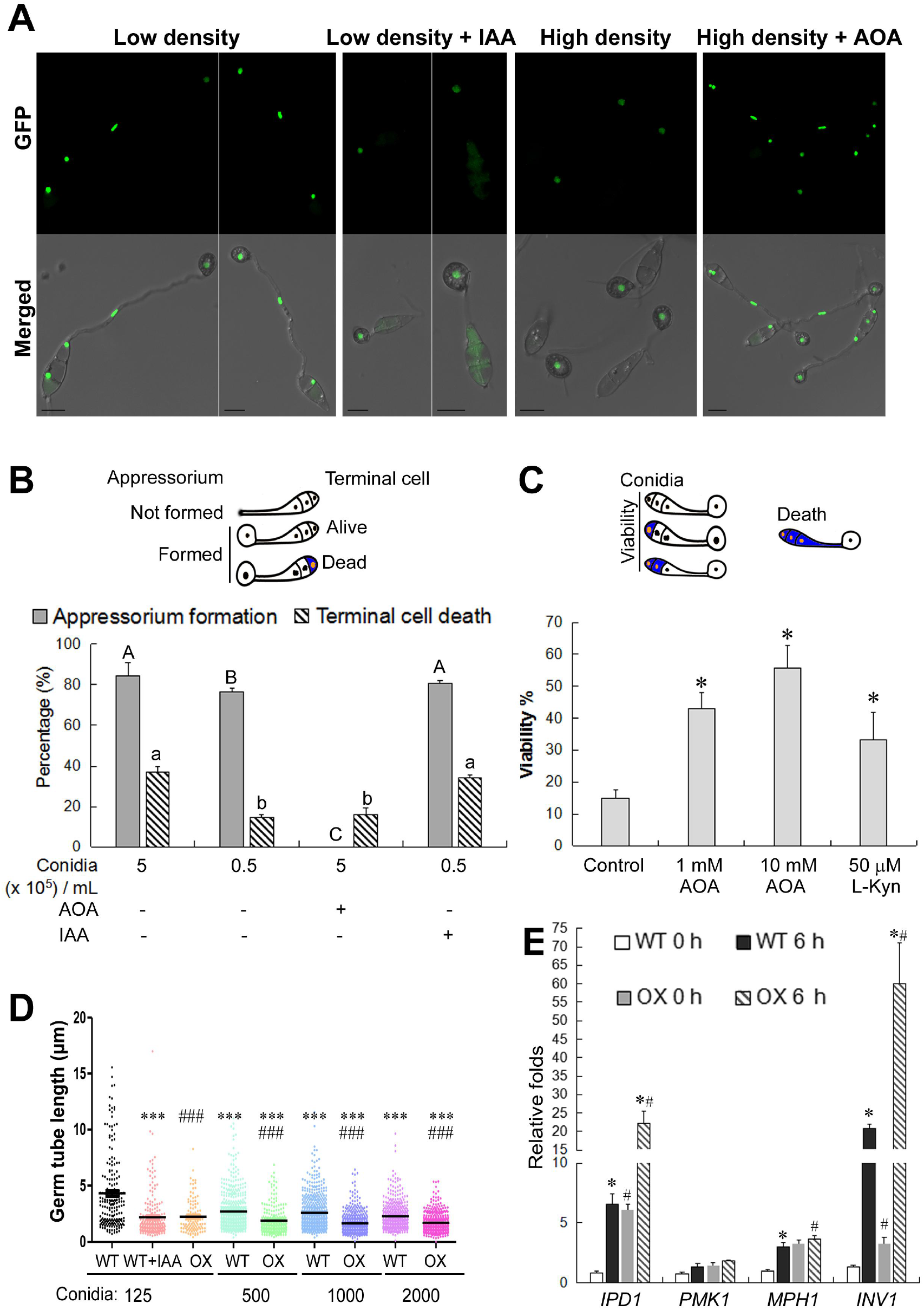
Auxin regulates *M. oryzae* pathogenic development in a quorum-sensing manner. (A) Cell-density dependent regulation of conidial cell death during appressorium development. The nucleus marking H1-GFP conidia of either low (0.5×10^5^ conidia/mL) or high (3×10^5^ conidia/mL) density were inoculated on the hydrophobic cover slips, and conidial cell death judging by the disappearance of H1-GFP marked nucleus was imaged at 15 hpi. 100 μM IAA or 1 mM AOA ( pH=7) was added at 0 hpi to the conidia of low density or high density, respectively. H1-GFP signal was observed using the Leica TCS SP8 X inverted microscope system, and is shown as maximum intensity projection. Bars = 10 μm. (B) Appressorium formation and death of the terminal conidial cell on a density/IAA level basis. Wild-type conidia of different inoculum with or without AOA (20 mM) or IAA (100 μM) were inoculated on the hydrophobic cover slips to induce appressorium formation. Percentage (%) of appressorium formation and terminal conidial cell death were quantified at 8 hpi. See the schematic cartons for quantification details. AOA was added to the conidia at 0 hpi or 4 hpi for quantification of appressorium formation or cell death, respectively. Quantification data is depicted as mean ± *SD* from at least 3 technical replicates (n = 100 each) in each instance.Different letters, *majuscule* for appressorium formation while *minuscule* for terminal cell death, denote significant difference (p<0.05). (C) Dose-dependent suppression of conidial cell death by Auxin inhibitor(s). Fresh conidia of H1-GFP (10^5^ conidia/mL) were inoculated on the inductive surface, and conidial cell viability was quantified at 24 hpi using trypan blue staining. 1 mM or 10 mM AOA, or 50 μM L-kyrurenine (L-Kyn) was added at 4 hpi. The schematic cartons illustrate quantification details. Barchart depicting quantification of conidial viability (mean ± *SD*) is derived from three independent repeats (n = 100 conidia each) for each treatment. * p < 0.05 versus control (water). (D) Germ tube length of Wild-type (WT) or *IPD1*-OX (OX) conidia of different inoculum in the presence or absence of IAA (50 μM). The germ tube length was measured at 8 hpi. Three independent biological repeats were performed, with n=300 conidia in each instance. ***: significant difference (p<0.001) vs. WT of 125 conidia per droplet; ###: significant difference (p<0.001) between OX and WT with the same number of conidia per droplet. (E) Gene expression in responses to changes of internal auxin levels. qRT-PCR analysis of gene expression of *IPD1, PMK1, MPG1* and *INV1* in the wild-type (WT) or *IPD1-OX* conidia at 0 hpi (ungerminated conidia) or inoculated on the hydrophobic cover slips for 6 hours (during appressorium formation). Relative folds were calculated using 2^-ΔΔCt^ method, with *TUBULIN* serving as an internal control. Gene expression levels in WT at 0 hpi were set as “1”. Mean ± *S.E*. is derived from three biological repeats, each containing three technical repeats. * significant difference (p<0.05) vs. 0 h of the same strain; # significant difference (p<0.05) between *IPD1*-OX and WT at the same time point.

To further understand the function of auxin/IAA as a QSM in regulating *M. oryzae* pathogenic development, we examined the expression of three selected infection-related genes: *MPG1* (encoding a hydrophobin) [28], *PMK1* [29], and *INV1* which encodes an invertase that functions as a potential QSM [9], in response to altered IAA levels using wild-type and IAA overproducing *IPD1*-OX strains at two pathogenic development stages, 0 and 6 h, that are differ in cellular IAA levels. The result showed that *IPD1* transcription was significantly induced in wild type at 6 hpi (Figure 3E), corresponding to more auxin/IAA accumulation at this time point (Figure 1A) as compared to 0 h. Interestingly, the infection-related gene *MPG1* and *INV1* were also significantly induced at 6 hpi in the wild type (Figure 3E), and they displayed higher expression in the IAA overproducing *IPD1*-OX strain at both 0 h and 6 h, as compared to those in wild type (Figure 3E), likely acting as a response to elevated IAA accumulation in the conidial population resulted from IAA secretion by the developing conidia (Figure 1C). In contrast, another infection related gene, *PMK1*, displayed no obvious transcriptional changes between the two time points or strains (Figure 3E), which in turn highlights the specificity of IAA regulated gene expression. Overall, these findings showed that fungal IAA production mediates pathogenic development through a quorum sensing mechanism.

### Ipd1 is a mitochondria-associated protein in *M. oryzae*

Next, we examined the subcellular localization of Ipd1-GFP under native regulation using the genetically complemented *ipd1*Δ strain (see Materials and Methods for details). The Ipd1-GFP signal appeared as cytosolic filaments and punctae in the mycelium (Figure 4A), which we inferred as mitochondria, and further verified by staining the mycelia with MitoTracker Red FM, to visualize the mitochondria. We found that the stained mitochondrial signal completely co-localized with Ipd1-GFP (Figure 4A). Therefore, we conclude that Ipd1 localizes to mitochondria in *M. oryzae*.

**Figure 4.**
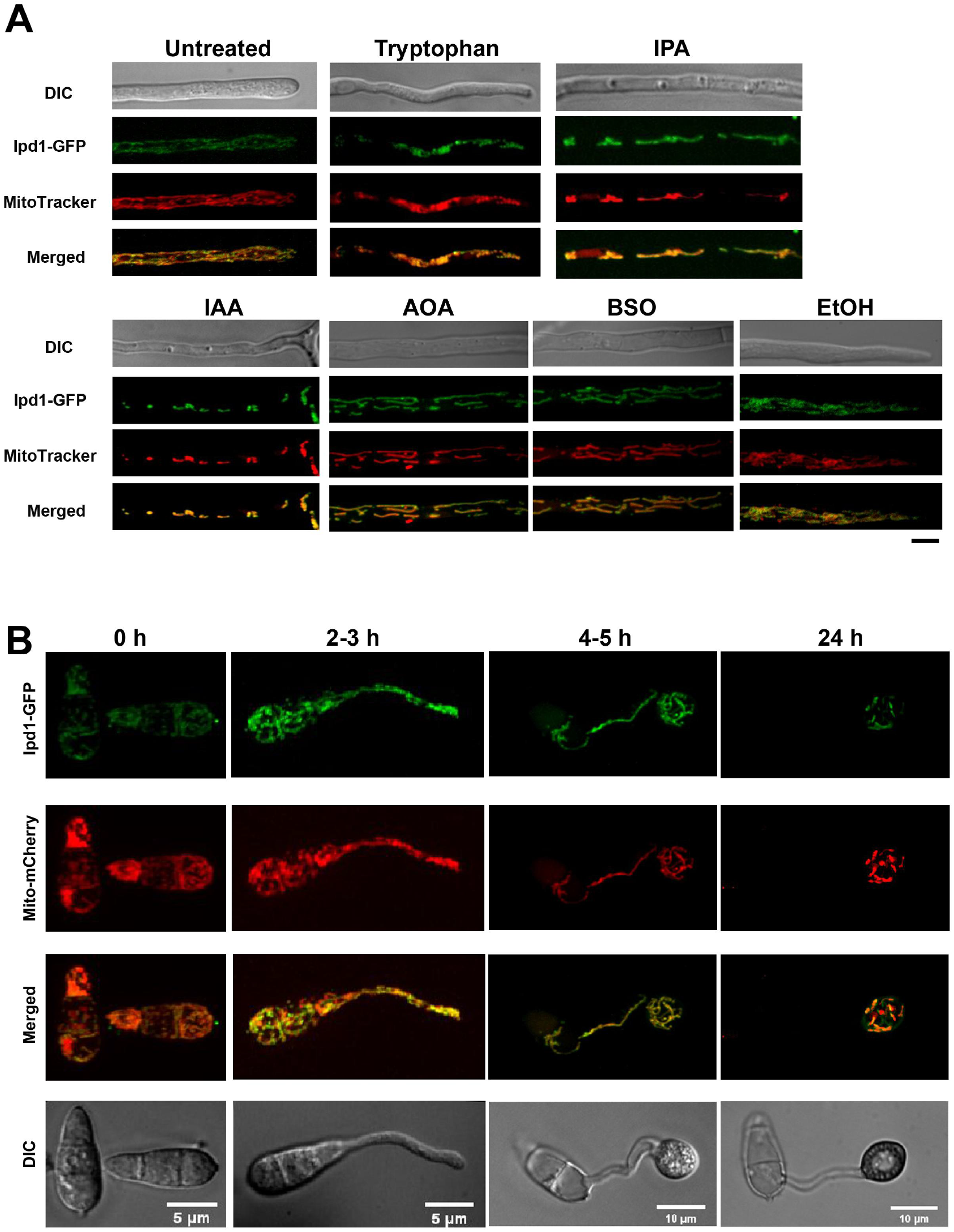
Mitochondrial localization of Ipd1-GFP during vegetative growth and pathogenic development. (A) Ipd1-GFP associated with mitochondria, stained by MitoTracker Red FM, under all conditions. Mycelia were grown in liquid CMN medium containing tryptophan (1 mg/mL), IPA (250 μM), IAA (50 μM), AOA (1 mM), BSO (100 μM) or ethanol (EtOH; 0.1%, solvent control) at 28 °C for 2 days. MitoTracker Red FM staining was performed at 28 °C for 30 min, and washed off with phosphatic buffer solution (PBS) for 30 min before epifluorenscent microscopy. Bar =5 μm. (B) Visualization of Ipd1-GFP co-localization with mitochondria. The conidia of Ipd1-GFP: Mito-mCherry co-expressing strain were inoculated on the hydrophobic cover slips. Ipd1-GFP fusion protein and Mito-mCherry were observed and imaged at the indicated time points. Images in this figure are all maximum intensity projections.

We further tested if Ipd1-GFP localization changes in response to treatment with IAA or precursor of IAA synthesis such as tryptophan and IPA. We noticed that tryptophan or IPA treatment caused increased/intensified Ipd1-GFP and MitoTracker Red signal (Figure 4A), indicating an enlargement of the filamentous mitochondrial network likely in response to a need for elevated IAA production induced by tryptophan or IPA (as substrates). In contrast, treatment with IAA led to reduced and predominantly punctate mitochondria (Figure 4A), likely due to the suppression of IAA production via negative feedback upon accumulation of the product. We also observed that treatment with AOA or applying oxidative stress using buthionine sulfoximine (BSO) did not change the subcellular localization of Ipd1-GFP (Figure 4A)

To observe the subcellular localization of Ipd1 during pathogenic development and host infection, we used the *IPD1*-OX strain (Figure S3E), in which the *IPD1-GFP* coding sequence was driven by the constitutive *RP27* promoter and the mitochondria were indicated by a stably expressed mitochondrial marker labeled as Mito-mCherry. Similarly, we found that the Ipd1-GFP fusion protein localized to the mitochondria in developing conidia and appressoria, although Ipd1-GFP was subsequently undetectable upon conidial cell death (Figure 4B). We also observed mitochondrial localization of Ipd1-GFP in the invasive hyphae, which was not affected by exogenous IAA addition (Figure S6C). Overall, we conclude that Ipd1 is a mitochondria-associated IPDC that catalyzes IAA production during *M. oryzae* pathogenic development and host infection.

In summary, our study demonstrates that the rice blast fungus produces phytohormone auxin during pathogenic development and host infection using the mitochondria-associated Ipd1. Such fungal auxin is secreted out and governs the severity of blast disease through a hitherto unknown quorum sensing mechanism. To our knowledge, this is the first report of fungus-sourced phytohormone serving a QSM function to regulate fungal pathogenesis.

## Discussion

Five tryptophan-dependent IAA synthesis pathways have been reported in plant and fungi, mostly based on the diverse intermediate products, namely IPA (Indole-3-Pyruvic Acid), IAM (Indole-3-AcetaMine), IAN (Indole-3-AcetoNitrile), TPA (TryPtAmine), and TSO (Tryptophan Side chain Oxidase) [22–23]. In this study, we identified an Indole-3 pyruvate decarboxylase Ipd1, and showed that it is crucial for *M. oryzae* IAA biosynthesis, as the *ipd1*Δ mutant produced significantly less IAA compared to the wild type, and the *IPD1*-overexpressed strain produced higher level of IAA (Figure S4). Correspondingly, the *ipd1*Δ mutant was defective in infecting the host rice (Figure S6A), while the *IPD1*-overexpressed strain displayed enhanced virulence and less sensitive to the IAA synthesis inhibitor AOA (Figure 2A). These results confirm that fungal IAA produced by Ipd1 contributes to fungal virulence.

Interestingly, such fungal IAA/auxin was secreted out (Figure 1C), accumulated in the fungal population on a cell-density basis (Figure 2C and Figure 3A, B, and D), functioned in a dose-dependent fashion (Figure 2A and D, and Figure 3C-D), and was able to trigger pathogenic responses from rice blast fungus (Figure 3D-E), which falls into the definition of quorum sensing. Quorum sensing was first reported and extensively investigated in bacteria, but its characterization was fairly limited in fungi/yeast, and mainly focused on dimorphic switching (yeast to pseudohyphal form) in *C. albicans, C. neoformans*, or in budding yeast [30]. In filamentous fungi *A. flavus* and *A. nidulans*, oxylipins and γ-Heptalactone have been identified as QSMs regulating fungal conidiation, virulence and/or secondary metabolism [31–32]. This study, for the first time, reports that a fungus-sourced phytohormone, auxin/IAA, serves a QSM function to regulate fungal pathogenesis, and may also act as an interkingdom signaling molecule to mediate the biotic interaction during rice and blast fungus.

By using the DII-Venus auxin biosensor in the fungus, we observed that auxin accumulated in the conidia in a sequential manner (from terminal conidial cell to the one proximal to the appressorium), in the process of appressorium development (Figure 1A). Such auxin accumulation pattern coincides with that of conidial ferroptosis, an iron-dependent cell death mediated by peroxidation of membrane lipids [33], during appressorium formation and maturation [20], and thus intrigued us to infer that auxin may regulate conidial ferroptosis. Indeed, our results showed that inhibition of IAA production by AOA effectively suppressed such conidial cell death (Figure 3 A-C). Unfortunately, we could not further verify this hypothesis by assessing conidial death in the *ipd1*Δ mutant, since this mutant fails to produce conidia. However, by searching the literature we found an established connections in plants between auxin and iron homeostasis [34–37], and between auxin and lipid oxidation [38–40], thus suggesting a potential relationship between fungal auxin dynamics and ferroptotic cell death in *M. oryzae*.

Recently, it is reported that auxin-secreting beneficial bacteria utilize auxin to counteract the plant immune response and ROS toxicity, thus facilitating host colonization by the bacteria, but meanwhile protecting the host plant against fungal pathogens likely via the activated plant immune response [41]. We imply that similar ROS detoxifying strategy may also be utilized by *M. oryzae* to suppress rice immunity during early host invasion stage. However, such auxin-secreting bacteria may not be applicable in protecting rice again blast disease, as our study demonstrated that auxin/IAA also plays a positive role in *M. oryzae* pathogenicity.

In summary, based on our results, we propose a working model for fungal auxin/IAA -mediated functions during pathogen-host interaction (Figure S7) in rice blast. The *M. oryzae* IPDC ortholog, Ipd1, is responsible for auxin/IAA production likely on an IPA-based pathway. Level of such fungal auxin/IAA correlates with the cell density, and determines the incidence and severity of blast disease in rice by affecting appressorium function, regulated conidial cell death, and infection-related genes expression. Elucidation of the upstream regulatory pathways and the downstream responsive elements of such intracellular auxin in *M. oryzae* will certainly help further our understanding of such novel fungal quorum sensing module in rice blast and other devastating pathosystems.

## Materials and Methods

### Strains, culture conditions and transformation

The *M. oryzae* strain B157 was used as wild type in this study. The wild type and the derived transformants/mutants were grown on prune agar medium (PA) or complete medium (CM) at 28 °C for 7 days. For detection of auxin in *M. oryzae*, or for examination of Ipd1-GFP subcellular localization, the fungal mycelia were grown in liquid complete medium with basal nitrogen (CMN) under dark condition (28 °C, 180 rpm) for 2 days. The colony sizes were measured at 6 days post inoculation (dpi). In this study, the deletion mutants and over-expression strains were generated by *Agrobacterium tumefaciens*-mediated transformation (ATMT) [42]. The genetic complementation strains were generated via protoplast transformation [43].

### Plant cultivar, growth, blast infection and auxin biosensor assays

Two-week-old seedlings of the rice cultivar CO39 and 7-day-old seedlings of the barley cultivar susceptible to *M. oryzae* were used for blast infection assays. The inoculated leaf explants were incubated in a growth chamber at 28 °C, 80% humidity, and a 12 h:12 h day:night cycle. The freshly harvested conidia at indicated concentrations, or the fungal mycelial plugs from 7-day-old cultures were used for the infection assays. Blast disease symptoms were examined and imaged at 7 dpi.

For fungal auxin detection by a plant biosensor, the freshly harvested wild-type *M. oryzae* conidia at a concentration of 10^6^ conidia/mL in sterile water were inoculated on cover slips (Menzel-Glaser) and extracellular fluid was collected at 6 h post inoculation (hpi). As control, same volume of sterile water without conidia was also inoculated on cover slips and collected in the same way. Extracellular fluid or water was then mix with Murashige and Skoog (MS) medium with agar (extracellular fluid: MS = 1: 1, v/v) in Petri dishes. Seven-day-old DR5:GFP seedlings (seeds purchased from Arabidopsis biological resource center, abrc.osu.edu/researchers; CS9361) germinated and grown on normal MS medium were transferred to the solidified half MS mixture and incubated for 30 min, and then stained with propidium iodide (PI; 10 μg/mL, Invitrogen P3566) at room temperature for 5-10 min to outline the viable *Arabidopsis* root.

### Chemical reagents used in this study

Amino-oxyacetic acid (AOA; Aldrich, C13408; pH adjusted to 7.0 when used); Buthionine sulfoximine (BSO; Sigma, B2515); Indole-3-acetic acid (IAA; Sigma, I2886); Indole-3-pyruvic acid (IPA; Sigma, I7017); L-kerurenine (L-Kyn; Sigma-Aldrich, K8625); Tryptophan (Trp, Sigma, T0254).

### Plasmid constructs and fungal transformants

The deletion construct pKO-IPD1, was generated based on the plasmid pFGL821 (with the hygromycin resistance gene, *HPH*) by flanking the resistant gene (selection marker) with the homologous regions of the targeting gene (Figure S3B). The deletion construct was transformed into the wild-type strain to generate the corresponding deletion mutants. For *IPD1* complementation, the *IPD1* locus, including its native promoter region (1.5 Kb), coding region (1.9 Kb, without stop codon) and the GFP coding sequence were PCR amplified and cloned into the vector pFGL932 to create the pIPD1:GFP construct, which was then transformed into the *ipd1*Δ protoplasts to generate complement strain.

To generate an *IPD1* over-expression strain, the *RP27* promoter, the coding sequence of *GFP* in-frame fused with *IPD1* gene, were PCR amplified and inserted into the plasmid pFGL1010 (Addgene, 119081, sulfonylurea resistance included) [44] sequentially to generate the pIPD1OX:GFP construct, which was then transformed into the wild-type strain. pMito-mCherry was constructed by inserting the promoter and first exon, which includes a mitochondria targeting sequence, of the gene encoding Enoyl-CoA hydratase (*ECH1*), and the mCherry coding sequence sequentially into the plasmid pFGL821. The resultant pMito-mCherry was transformed into the above-mentioned *IPD1-GFP* over-expression strain.

To generate the auxin reporter DII-Venus in *M. oryzae*, the *RP27* promoter and the DII-Venus-NLS fusion sequence [18] were PCR amplified and inserted into pFGL1010 to obtain the p*RP27*-DII-Venus construct, which was then transformed into the wild-type strain by ATMT. The pFGL1170R (Addgene, 116896) containing a nuclear marker H1-mCherry was then transformed into the above-mentioned DII-Venus strain by ATMT to get a DII-Venus and H1-mCherry co-expressing strain. The primers for plasmid construction and for mutant verification are listed in Table S2.

### Nucleus acid manipulation

Total RNA was extracted from the mycelia using RNeasy Plant Mini kit (QIAGEN, United States). The first strand cDNA synthesis and qRT-PCR were carried out as mentioned [45]. The primers used for qRT-PCR are listed in Table S3. The genomic DNA was extracted from mycelia using SDS protocol [46]. Southern blot analysis was performed following an established protocol [47].

### IAA detection using liquid chromatography mass spectrometry (LC-MS)

Sample preparation: The liquid cultured mycelia were ground to a fine powder using liquid nitrogen and a mortar pestle. The resultant powder was mixed with auxin extraction buffer (isopropanol: water: HCl=2:1:0.001, v/v) for 4-5h using an oscillator under dark conditions. The mixture was further mixed with dichloromethane (Sigma, D65100) for another 2-3h. For every 100 mg power, 600 μL auxin extraction buffer and 600 μL dichloromethane were used. The final mixture was then centrifuged at 12,000 rpm for 10 min, and the precipitate was then dried and re-dissolved in 400 μL 50% methanol. For exacting auxin from the medium, mycelia were removed via filtration and the left medium was mixed with equal volume of ethyl acetate (pH=2.0, adjusted using HCl) on the oscillator for 4-5h under dark conditions. The mixture was then centrifuged at 12,000 rpm for 10 min, and the supernatant was taken, dried and re-dissolved in methanol (1 mL methanol was used for 10 mL medium). The solution was diluted 100 times with 50% methanol.

#### LC-MS

The analysis was carried out in an ultra-high performance liquid chromatography (UHPLC) system (Agilent 1290 Infinity, USA) coupled to LCMS (Agilent 6490 Series Triple Quadrupole, USA), controlled by MassHunter software B.06.00. The UHPLC system was equipped with a Kinetex C18 column (100 × 2.1mm, 1.7 μm, 100 Å, Phenomenex) heated at 50 °C. For auxin analysis, 10 μL of extracts were injected and followed by the separation at a constant flow rate of 0.3 mL/min, in a gradient of solvent A (water acidified with 0.1% formic acid) and B (acetonitrile acidified with 0.1% formic acid): 1 min 5% solvent B; 9.5 min 5% to 100% solvent B; 2.9 min 100% solvent B, and re-equilibration to the initial conditions in 3 min. The setting for the mass spectrometer conditions were as follows: Electrospray Ionization (ESI) source temperature 250°C, electrospray voltage at −4000V (negative mode), gas flow at 12 L/min, nebulizer gas pressure 35 psi, sheath gas temperature 350°C, sheath gas flow 11 L/min. Certified auxin standard was purchased from Sigma (I2886). Standard stock solution (1,000 μg/mL) was prepared in the methanol as solvent.

### Microscopy, image analysis and processing

Staining with MitoTracker Red FM (Molecular probes, M22425) was carried out as described [10]. The fluorescence microscopy was performed using an UltraView RS-3 spinning disk confocal system (PerkinElmer Inc., United States), with a 491 nm 100 mW and a 561 nm 50 mW laser illumination under the control of MetaMorph Premier Software. Image processing was carried out using ImageJ (version 1.8.0-172), Adobe Photoshop (2017 version) and Adobe Illustrator (2017 version).

Epifluorescence of DR5:GFP, PI, and H1-GFP (for cell viability/cell death) was observed using a Leica TCS SP8 X inverted microscope system (Leica Microsystems) equipped with an HC Plan Apochromat 20×/0.75 CS2 Dry objective or an HC Plan Apochromat 63×/1.40 CS2 Oil objective, respectively. The white light laser controlled by the AOTF (Acousto-Optical-Tunable-filter) for rapid modulation of intensity was used for GFP (excitation, 488 nm; emission, 500-530 nm) and PI (excitation, 561 nm; emission, 600-700 nm). All the images were captured using the Leica Hybrid Detector. All parts of the system were under the control of Leica Application Suite X software package (release version 3.5.5.19976).

## Data analysis

For LC-MS/MS analysis, Qualitative Analysis (version B.06.00), MultiQuant (version 3.02) or Analyst (version TF1.17) was used for quantitative data processing. For statistical analysis, the one-way analysis of variance (ANOVA) tests was carried out (p <0.05, significant).

## Acknowledgements

We thank Dr. Zheng Wenhui for help with confocal microscopy, and the Shanghai Applied Protein Technology Co. Ltd; and NUS Environmental Research Institute, Singapore for technical support in metabolomics analysis. We thank the members of Deng group (SCAU, China) and Fungal Pathobiology Group (TLL, Singapore) for helpful discussions and suggestions.

## Author Contributions

YZD and NIN conceived and designed the study, and provided materials and funding support. LD, QS, CyC and LS performed the experiments. FY constructed the vectors used in this study. LD, QS, YZD and NIN analyzed the data and drafted the manuscript. All the authors reviewed and approved the manuscript.

