## supplemental Files for "Fungal auxin is a quorum-based modulator of blast disease severity"

#### Figure S1

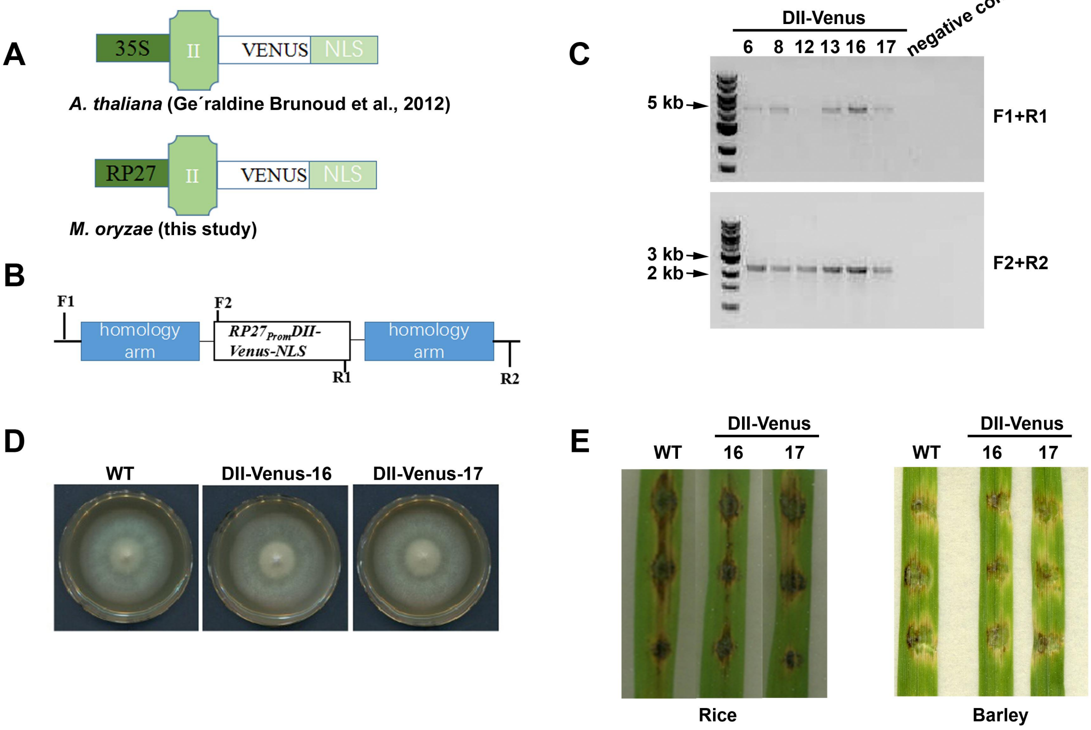


**Figure S1. Generation of auxin/IAA reporter strain.** (A) Design of DII-Venus in *M. oryzae* on the basis of DII-Venus in *A. thaliana* [1]. The fusion protein was expressed under the strong *RP27* promoter. (B and C) The DII-Venus strains were verified by PCR amplification (C) with two primer pairs (B, sequences listed in Table S2). (D) Similar radial growth of the wild-type (WT) and the DII-Venus: H1-mCherry strains (#16 and #17) on PA medium assessed at day 6. (E) No obvious difference between wild type and DII-Venus: H1-mCherry strains in causing blast disease on rice or barley leaves. The conidial suspension droplets (each contains 2,000 conidia) from wild type (WT), DII-Venus: H1-mCherry strain #16 or #17 were placed on rice or barley leaf explants, and incubated for one week before photographing.

### Figure S2

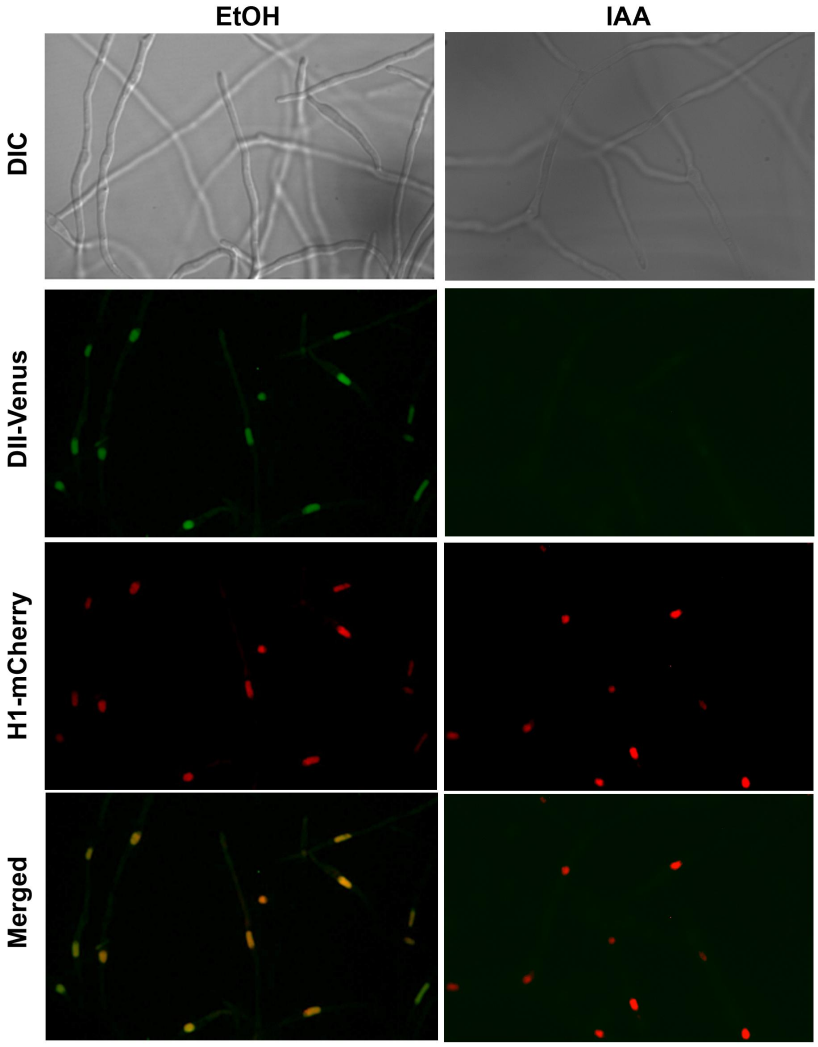


**Figure S2. IAA triggers the degradation of engineered DII-Venus.** The vegetative mycelium of DII-Venus: H1-mCherry strain (#16 as a representative) was grown in liquid CMN with or without 50 μM IAA for 2 days under dark condition. 0.1% ethanol (EtOH) served as solvent control. The DII-Venus and H1-mCherry signal were observed using a spinning disk confocal microscope. Single plane images are shown here. Bar = 5 μm.

### Figure S3

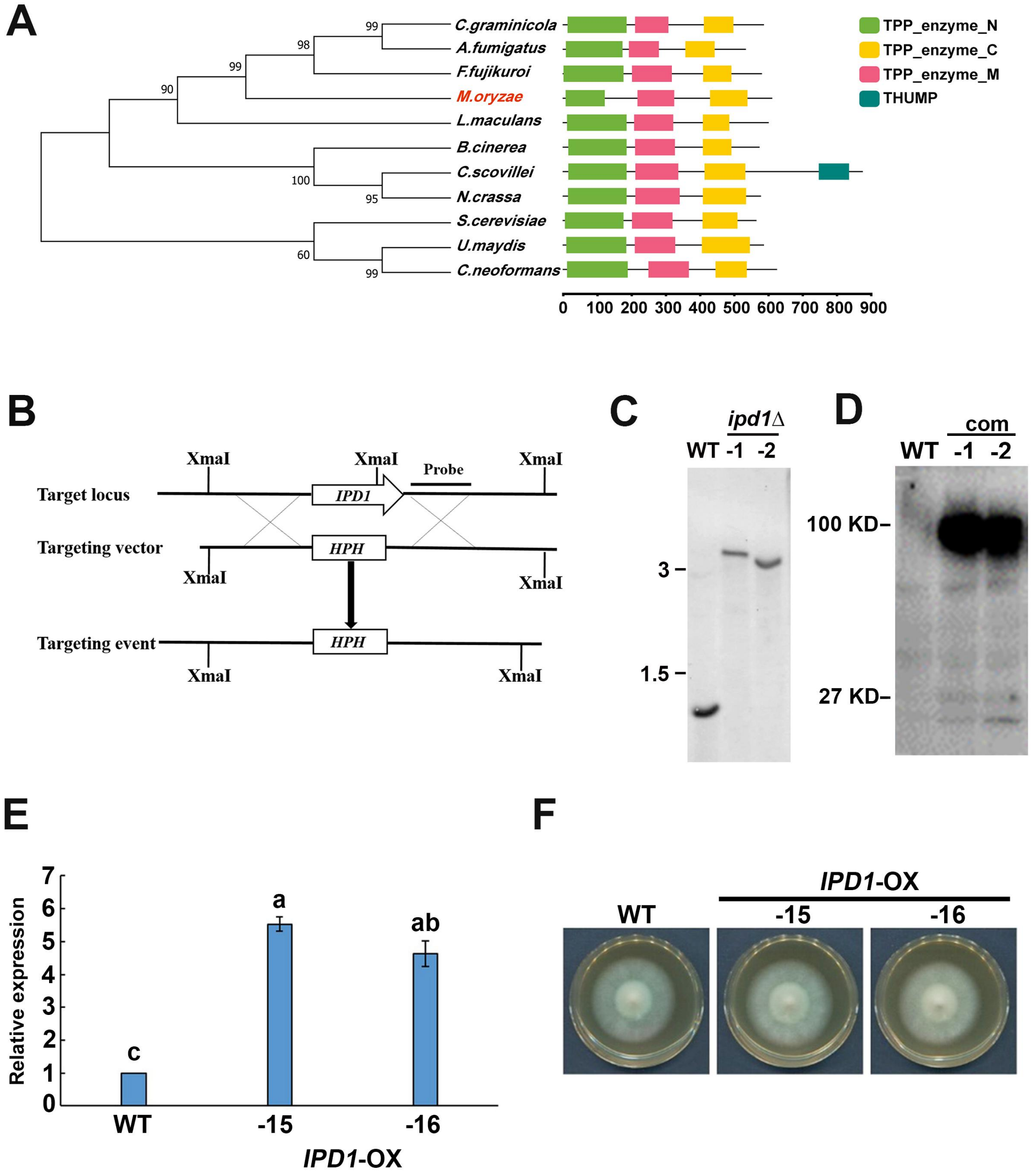


**Figure S3. Phylogenetic analysis of Ipd1 orthologs and generation of *IPD1* deletion, complementation, and over-expression strains. (A)** The evolutionary history was inferred using the Neighbor-Joining method [2]. The percentage of replicate trees in which the associated taxa clustered together in the bootstrap test (1,000 replicates) were shown next to the branches [3]. Branches corresponding to partitions reproduced in less than 50% bootstrap replicates were collapsed. The evolutionary distances were computed using the Poisson correction method [4] and in the units of the number of amino acid substitutions per sites. Evolutionary analysis were conducted using MEGA7 [5]. Domain annotation is drew to scale using TBtools software [6], and the numbers under the scale denote amino acid resides. (B) Schematic representation showing how *IPD1* deletion was performed. The *IPD1* gene was replaced by a hygromycin phosphotransferase cassette (*HPH*). The sites of restriction enzyme digestion and the site used to design probe for southern blot are marked. The schemes are not drew to scale. (C) Southern blot analysis of *IPD1* deletion mutants. Probe (see details in (B)) detected a single band of 1.2 kb in the wild type (WT), but a single band of 3.6 kb in the *ipd1*Δ mutants. (D) Verification of complementation strains by detecting a GFP-Ipd1 fusion protein (approximately 94 kDa) using anti-GFP antibody via immunoblotting. Total lysate from the wild-type (WT) mycelia served as negative control. (E) *IPD1* over-expression verified by qRT-PCR. Relative expression was calculated using $2^{-\Delta\Delta Ct}$ method using *TUBULIN* as a internal control. Gene expression level in wild type (WT) was set as "1". Mean ± *S.E.* based on three independent biological repeats is shown. Different letters indicated significant differences ($p < 0.05$). (F) Vegetative growth of *IPD1* over-expression strains (-15 and -16) was assessed after grown on PA medium for 6 days.

### Figure S4

A

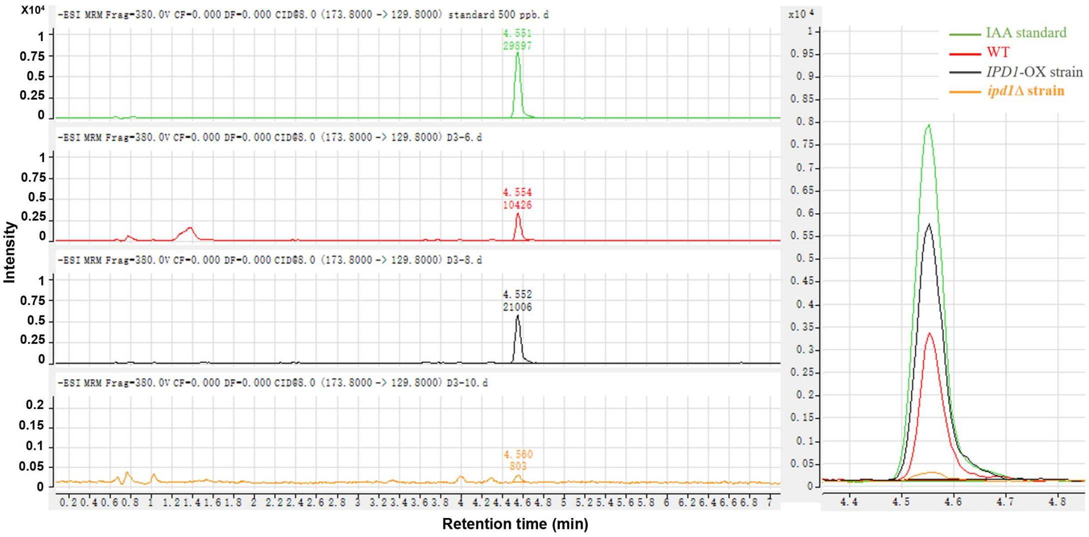


B

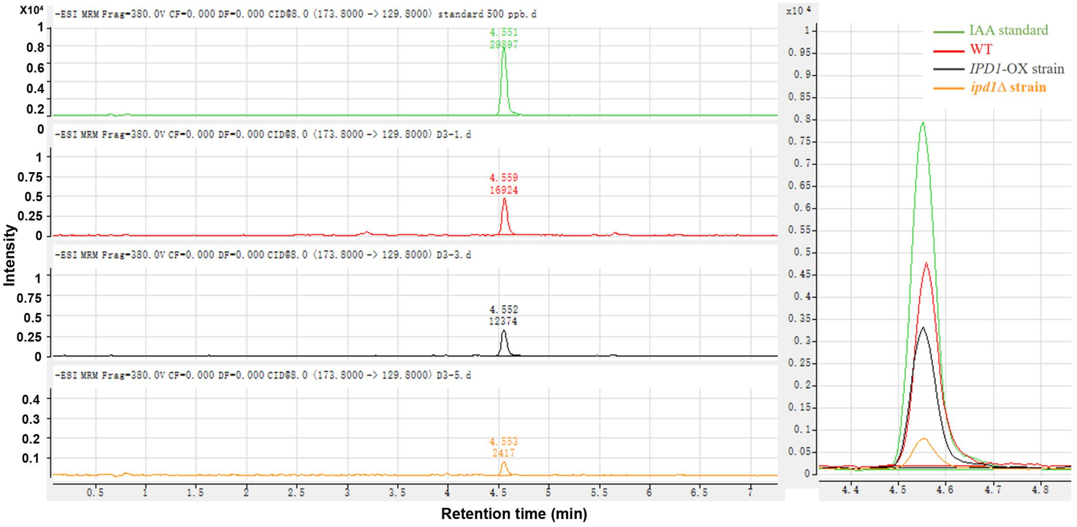


**Figure S4. Ipd1 is essential for auxin/IAA biosynthesis in *M. oryzae*.** The mycelia of wild type (WT), *ipd1*Δ mutant or *IPD1* over-expression (OX) strain were cultured in liquid CMN medium with 1 mg/mL tryptophan under dark condition for 2 days. Auxin/IAA production either within fungal cells (A) or secreted out (B) was reduced in the *ipd1*Δ mutant compared to wild type. Increased auxin/IAA production within *IPD1*-OX strain was detected (A). IAA (Sigma, I2886) of 500 ppb served as a standard.

### Figure S5

A

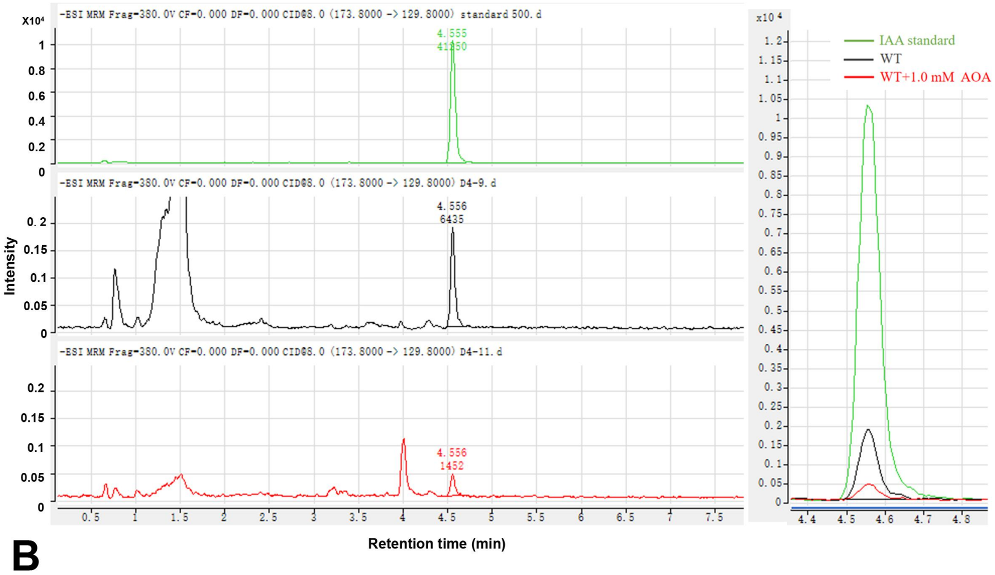


B

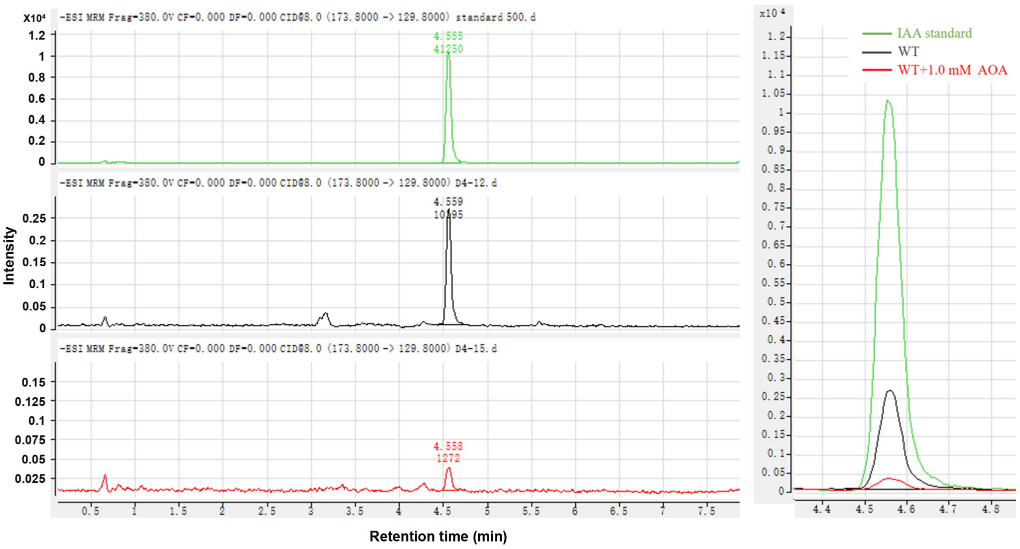


**Figure S5. AOA suppresses both intracellular and secreted auxin levels.** Wild-type (WT) mycelia were allowed to grow in CMN liquid medium containing 1 mg/mL tryptophan, under dark condition for 2 days. Treatment by 1 mM AOA caused obvious reduction of IAA production either within (A) or outside (B) the fungal cells. IAA of 500 ppb was used as a standard.

### Figure S6

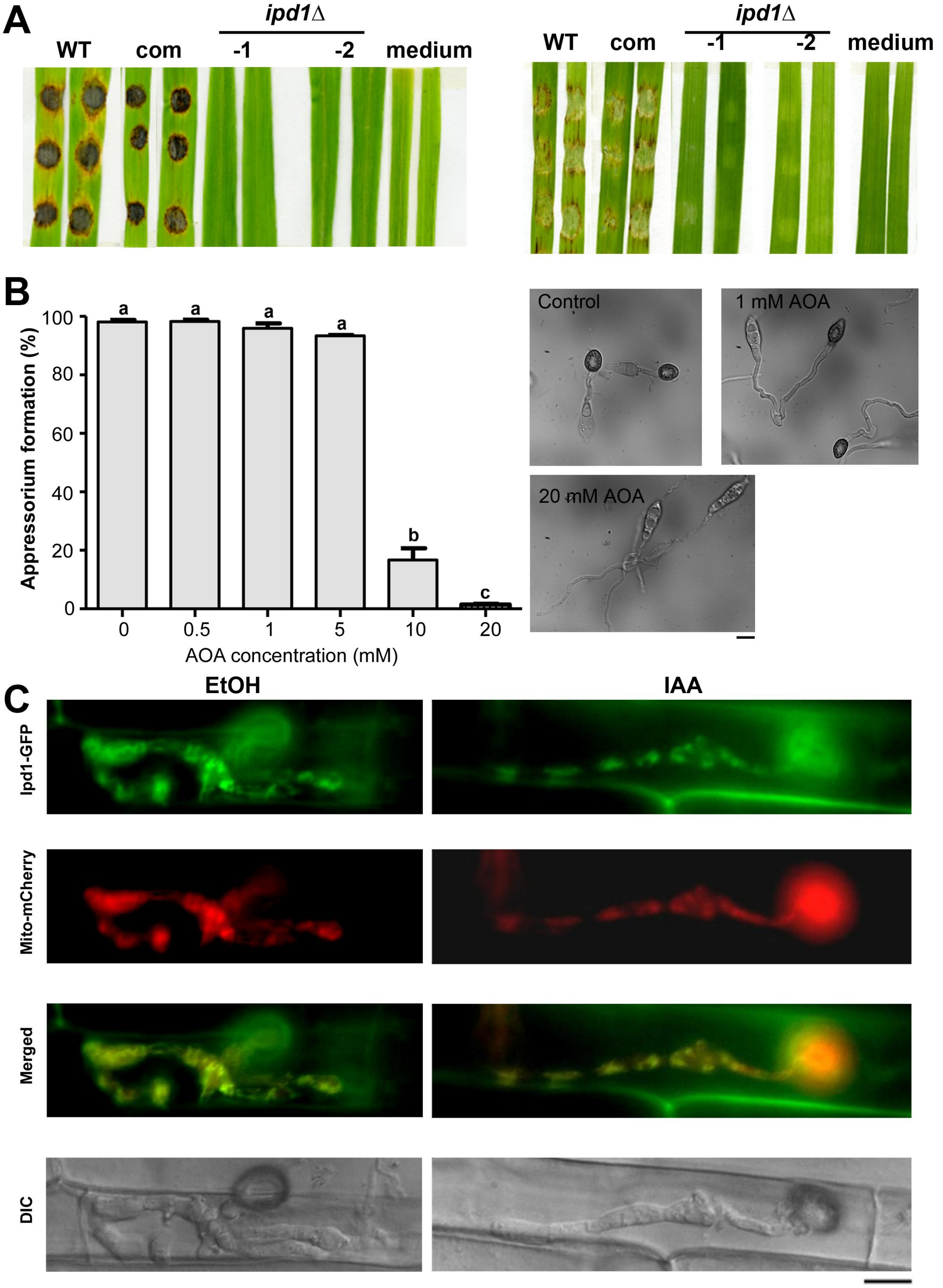


**Figure S6. Mitochondria associated Ipd1 is essential for *M.oryzae* pathogenicity.** (A) Infection assay using two-week-old rice (left panel) or one-week-old barley (right panel) explants. The mycelial plugs of wild type (WT), *ipd1*Δ mutants, or complementation strain (com) was inoculated on the leaf explants. Fresh medium plugs served as blank controls. Photos were taken at 7 dpi. (B) Delay or suppression of appressorium formation by low and high AOA dosages, respectively. The fresh wild-type conidia were inoculated on the inductive surface. AOA of different concentrations, was added to the conidial suspension ($10^5$ conidia/mL) at 0 hpi, and the percentage (%) of appressorium formation was quantified at 8 hpi. Mean ± *S.D.* is derived from three independent repeats (n = 100 conidia each) for each treatment. Different letters denote statistical significance ($p < 0.05$) versus control (water). Representative DIC images are showed. Bar = 10 μm. (C) The conidia of Ipd1-GFP: Mito-mCherry co-expressing strain were inoculated on the rice leaf sheath, with or without supplement of 50 μM IAA. 0.1% ethanol (EtOH) served as solvent control. Ipd1-GFP and Mito-mCherry were observed and imaged at 30 hpi. Images shown are maximum intensity projections. Bar = 10 μm.

### Figure S7

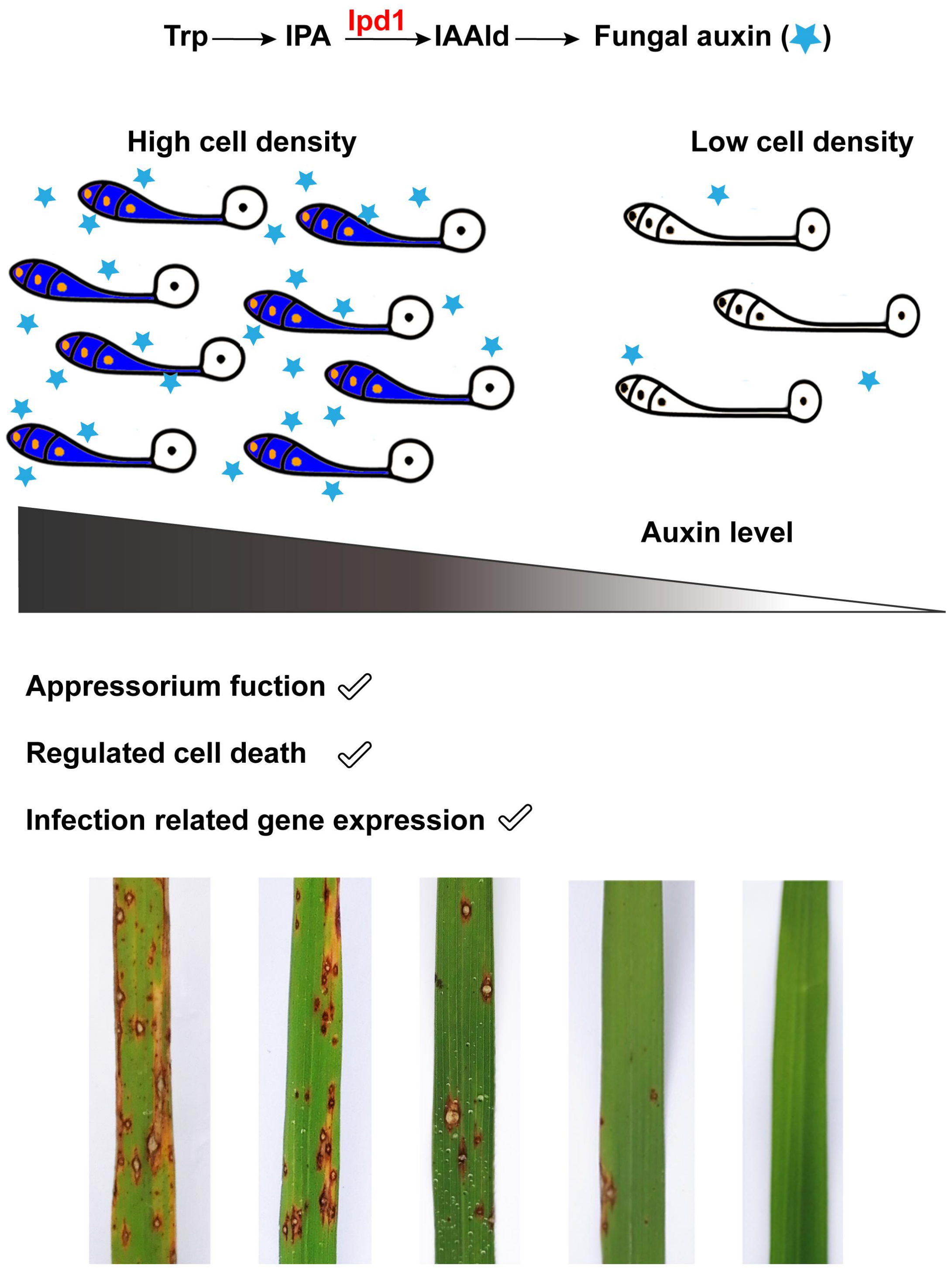


**Figure S7. Proposed working model for fungal auxin/IAA -mediated blast-rice interaction.** *M. oryzae* Ipd1 is responsible for auxin/IAA production likely on an Indole-3- Pyruvic Acid (IPA)-based pathway using Tryptophan (Trp) as the substrate. Level of such fungal IAA correlates with the conidial cell density, and determines the appressorium function, regulated cell death occurring in the conidium, and expression of infection-related genes on a quorum-sensing basis, therefore determines the incidence and severity of blast disease in rice.

**Table S1. Comparison of mycological characteristics among strains.**

| **Strain** | **Radius of colony (average ± *S.E.*)** | **Conidiation count (×$10^5$/mL; average ± *S.E.*)** |
|---|---|---|
| Wild type (WT) | 3.52±0.02[a] | 58.76±5.08[a] |
| *ipd1*Δ-1 | 1.78±0.02[b] | 0[b] |
| *ipd1*Δ-2 | 1.80±0.03[b] | 0[b] |
| *IPD1*-com-1 | 3.51±0.02[a] | 55.98±5.32[a] |
| *IPD1*-com-2 | 3.56±0.03[a] | 60.12±3.13[a] |
| *IPD1*-OX-15 | 3.68±0.02[a] | 49.63±4.21[a] |
| *IPD1*-OX-16 | 3.74±0.02[a] | 55.06±1.91[a] |
| *IPD1*-OX: mito-mCherry-1 | 3.71±0.03[a] | 55.73±1.81[a] |
| *IPD1*-OX: mito-mCherry-2 | 3.75±0.01[a] | 54.05±1.32[a] |
| DII-Venus:H1-mCherry-16 | 3.69±0.01[a] | 55.06±3.80[a] |
| DII-Venus:H1-mCherry-17 | 3.75±0.03[a] | 56.82±3.01[a] |

Difference letters denote statistic significance ($p<0.05$)

**Table S2. Primers used for plasmid construction.**

| Gene (Locus) | Description | Enzyme sites | Primer sequence |
|---|---|---|---|
| *IPD1* (*MGG_01892*) | Deletion construct | *Eco*RI | UPF: 5'-CCG**GAATTC**GCTGTTGCCATTGAATCC-3' |
| | | *Xba*I | UPR: 5'-TGC**TCTAGA**CGGTGGTTTGGTCAAGGTT-3' |
| | | *Pst*I | DSF: 5'-AA**CTGCAG**GCTTTCACGCCAGGTATGA-3' |
| | | *Hind*III | DSR: 5'-CCC**AAGCTT**GATTCCAGGTTCGTGTTTG-3' |
| | PCR verification | — | 5'-CCCTTTATCCCATTCAAG-3' |
| | | — | 5'-AGGCATCCATTCCAAGACC-3' |
| | | — | 5'-CAATGTCAAGCACTTCCG-3' |
| | Complementation | *Nco*I | IPD1F:5'-CATG**CCATGG**CCCTTTATCCCATTCAAGAC-3' |
| | | — | IPD1R: 5'-GCCCTTGCTCACCATTTCTGGATCACTCCGGATCC-3' |
| | | — | GFPF: 5'-CGGAGTGATCCAGAAATGGTGAGCAAGGGCGAGGAGC-3' |
| | | *Eco*RI | GFPR: 5'-CCG**GAATTC**TTACTTGTACAGCTCGTCCATGCCG-3' |
| | Overexpression | *Xho*I | RP27F: 5'-CCG**CTCGAG**ATAAATGTAGGTATTACC-3' |
| | | *Eco*RI | RP27R: 5'-CG**GAATTC**TTTGAAGATTGGGTTCC-3' |
| | | *Eco*RI | IPD1F: 5'-CG**GAATTC**ATGACCAAGATCCCCCTCG-3' |
| | | *Kpn*I | IPD1R: 5'-GG**GGTACC**TTCTGGATCACTCCGGAT-3' |
| | Subcellular localization | *Eco*RI | 5'-CG**GAATTC**TCGATAATCTGGATGTCTGC-3' |
| | | *Xba*I | 5'-TGC**TCTAGA**AAGAAGGATTACCTCTA-3' |
| *DII-Venus-NLS* | Auxin reporter construct | *Eco*RI | RP27F: 5'-CATGATGATGCTCGA**GAATTC**ATAAATGTAGGTATTACC-3' |
| | | *Kpn*I | RP27R: 5'-ACTAGTGGATCCCCG**GGTACC**TTTGAAGATTGGGTTCC-3' |
| | | *Xba*I | 5'-CGGGGATCCACTAGT**TCTAGA**ATGAAACAAAAAAGCTCGACCAAAG-3' |
| | | *Pst*I | 5'-CCCTGTGGAGCATGC**CTGCAG**TTACTCTTCTTCTTGATCAGCTT-3' |

**Table S2 continued**

| Gene (Locus) | Description | Enzyme sites | Primer sequence |
|---|---|---|---|
| | PCR verification | — | F1: 5'- GATTGTGTGGAGGGACTTG-3' |
| | | — | R1: 5'- TTACTCTTCTTCTTGATCAGCTT-3' |
| | | — | F2: 5'- ATGAAACAAAAAAGCTCGACCAAAG-3' |
| | | — | R2: 5'- CCTTGTGTGGTGTCGTGAAG-3' |

**Table S3. Primers used for qRT-PCR.**

| Gene (Locus) | Primer sequence |
|---|---|
| *PMK1* | F: 5'-CAATCCACCAAGCAACTC-3' |
| (*MGG_09565*) | R: 5'-TCGTAGGCAGAACATAGAAT-3' |
| *MPG1* | F: 5'-GAAGGTCGTCTCTTGCTGCA-3' |
| (*MGG_10315*) | R: 5'-GGATGTTGACCAGACCAATC-3' |
| *IPD1* | F: 5'-CGAGGAGTACAACGACAT-3' |
| (*MGG_01892*) | R: 5'-TGAACCTGCCATCCATAA-3' |
| *INV1* | F: 5'-GACTGGCTGCTGAATAAG-3' |
| *(MGG_05785)* | R: 5'-ATTGGTAGTGAAGGTGGTA-3' |
| *TUBULIN* | F: 5'-GTTCACCTTCAGACCGG-3' |
| (*MGG_00604*) | R: 5'-GAGATCGACGAGGACAG-3' |
